# A novel Gorilla-derived oncolytic Adenovirus with natural selective replication in cancer cells

**DOI:** 10.64898/2026.02.26.708271

**Authors:** Romina Scala, Ilaria Cela, Emily Capone, Valentina Proganò, Angiolo Pierantoni, Stefano Colloca, Gianluca Sala, Angelo Raggioli

## Abstract

Oncolytic virotherapy exploits viruses to selectively infect and destroy cancer cells while sparing normal tissues and represents a promising strategy in oncology. Human adenovirus type 5 (HAd5), although widely used, shows limited clinical efficacy due to high levels of preexisting immunity and suboptimal tumor selectivity. In this study, we evaluated novel gorilla-derived adenoviruses (GRAd) as alternative oncolytic vectors. Two distinct GRAd groups, GRAdBs and GRAdCs, were characterized for replication and cytopathic activity. GRAd25 (GRAdB group) exhibited robust replication in both tumor and normal cells, whereas GRAd32 (GRAdC group) demonstrated selective replication in tumor cells.

To broaden tumor tropism while preserving selectivity, we generated a chimeric GRAd32 vector, GRAd32Fk25, by replacing its native fiber knob with that of GRAd25, potentially shifting receptor usage from CAR to CD46, which is more abundantly expressed in tumor cells. The vector was further armed with a therapeutic antibody by inserting the coding sequence for the single-chain Fc form (scFv-Fc) of EV20, a humanized anti-HER3 antibody, under endogenous viral regulatory control.

In vitro analyses showed that GRAd32Fk25 maintained tumor-restricted replication and produced functional EV20 capable of binding HER3 and inhibiting downstream PI3K/Akt signaling. These results indicate that engineered GRAd vectors, exemplified by GRAd32Fk25 armed with EV20, provide a selective and versatile platform for oncolytic virotherapy with potential advantages over HAd5-based approaches.

## Introduction

Oncolytic virotherapy is an innovative approach in cancer treatment that employs viruses engineered or selected to preferentially infect and destroy tumor cells while sparing normal tissues. These oncolytic viruses (OVs) exert their therapeutic effects through two key mechanisms: direct tumor cell lysis and immune system activation. By inducing immunogenic cell death (ICD), OVs release tumor-associated antigens (TAAs) along with immune-stimulatory signals, including damage-associated molecular patterns (DAMPs), pathogen-associated molecular patterns (PAMPs), and pro-inflammatory cytokines, thereby priming an antitumor immune response that extends beyond infected cells^1^. This therapeutic strategy has led to the approval of several oncolytic viruses, including ONYX-015 (Oncorine), the first oncolytic adenovirus approved for head and neck cancer in China^2^, Rigvir, a reovirus approved for melanoma in Eastern Europe^3^, T-Vec (IMLYGIC), a GM-CSF expressing herpes simplex virus approved for melanoma^4^ in the Unites Stated, Europe and Australia, and Deltyact (Teserpatricev/G47Δ), a genetically modified herpes virus approved in Japan for glioblastoma^5^. With over 260 OV-based clinical trials initiated worldwide to date (www.clinicaltrials.gov, accessed December 15, 2025; keyword: “oncolytic virus”) the clinical landscape of oncolytic virotherapy remains dominated by a limited number of viral platforms and is largely driven by platform-centered rather than tumor-centered development strategies. Within this context, adenoviruses have assumed a central role, with the vast majority of clinical programs relying on adenovirus serotype 5 (Ad5) or Ad5-derived backbones. These include early-generation vectors such as H101^2,6^ and CG0070^7^, as well as biologically selective replication-competent adenoviruses based on E1-modified or fiber-modified Ad5 architectures, which have been repeatedly repurposed across multiple tumor indications^8,9^. Notably, Enadenotucirev (ColoAd1)^10^, a chimeric human group B Ad11p/Ad3 virus generated by directed evolution, represents a rare and significant departure from the Ad5 paradigm and remains the only non-Ad5 adenoviral oncolytic platform to have progressed extensively into clinical development. While this concentration around Ad5-based vectors - extending also to their widespread use as gene therapy and vaccine platforms - underscores their manufacturability and engineering flexibility, it has also exposed intrinsic limitations of human adenoviruses, including high seroprevalence^11–14^, off-target sequestration, and restricted biodistribution following systemic administration^15^. These limitations have motivated the exploration of alternative adenoviral species with distinct biological and immunological properties^16–19^. Among those, notable examples in the oncolytic field include the gorilla-derived group B oncolytic adenovirus GoraVir^20,21^, primarily evaluated in preclinical models of pancreatic ductal adenocarcinoma, and the chimpanzee-derived group E oncolytic adenovirus C7^22^. Together with ColoAd1, these vectors represent non-group C adenoviral alternatives to the canonical group C Ad5 backbone, that has historically served as the predominant platform for oncolytic and gene therapy vectors, and remains a benchmark for immunological potency^23^.

While diverging from group C can confer advantages such as reduced preexisting neutralizing immunity and additional distinct biological properties, it may also entail trade-offs in immunological potency. To overcome these challenges, we explored our collection of gorilla-derived adenoviruses (GRAd) and identified a unique gorilla-derived group C vector with the unexpected ability to replicate selectively in cancer cells while sparing normal tissues. Unlike HAd5, which requires extensive genetic modifications to achieve tumor selectivity^8,24–26^, this GRAd vector naturally exhibits cancer-specific replication, eliminating the need for complex genetic alterations that can limit therapeutic applicability.

HER-3 (ErbB-3), a kinase-impaired member of the EGFR family, plays a central role in mediating resistance to targeted therapies across several malignancies through activation of the PI3K/Akt pathway. Its expression is frequently upregulated in cancers such as melanoma, breast, ovarian, and pancreatic tumors, particularly under selective pressure from treatments targeting EGFR or HER-2. Several HER-3 targeting antibodies are currently in different stages of clinical evaluation, both as monotherapies and in combination with other anticancer treatments, for a range of solid tumors^27–30^. EV20 is a humanized monoclonal antibody specifically designed to target HER-3 with high affinity and clinical suitability^31^. Derived from the murine precursor MP-RM-1, EV20 was engineered to retain antigen specificity while minimizing immunogenicity and enhancing internalization into tumor cells^32^. EV20 antibody is demonstrated to be able to disrupt HER-3 signaling by inducing receptor downregulation and blocking both ligand-dependent and independent activation. In preclinical models *in vivo*, EV20 has demonstrated significant antitumor activity across multiple tumor types^31^. Beyond its use as a standalone agent, EV20 has been conjugated to cytotoxic payloads such as plant toxin Saporin, monomethyl auristatin F (MMAF) and duocarmycin derivative (NMS-P945) forming antibody-drug conjugates (ADCs) with enhanced efficacy and reduced systemic toxicity^33–37^. These EV20-based ADCs exhibit potent, selective therapeutic activity in HER-3 expressing solid tumors, including models of therapeutic resistance^30^. Importantly, EV20 has shown strong synergistic potential in combination therapies. Notably, its co-administration with Osimertinib, a third-generation EGFR inhibitor used in resistant non-small-cell lung cancer (NSCLC), exploits compensatory HER-3 upregulation to amplify therapeutic effects. This combination not only augments tumor cell killing but also activates innate immune responses, including macrophage engagement and STING pathway activation, thereby enhancing the overall antitumor efficacy by bridging targeted therapy and immunotherapy^38^.

Previously, a HAd5-based oncolytic adenovirus expressing the monoclonal antibody trastuzumab has shown promising *in situ* antibody production, coupled with progressive oncolytic cancer cell killing^39^. Here, we explored the incorporation of single-chain Fc (scFv-Fc) derivative of EV20 into the GRAd genome as a strategy to enhance therapeutic efficacy. This strategy is designed to allow the oncolytic virus to function as a biofactory for *in situ* monoclonal antibody production, thereby confining EV20 activity to tumor cells that support active GRAd replication. Such localized antibody delivery is expected to modulate downstream HER-3 signaling in malignant cells and, together with virus-induced oncolysis, promote immune cell activation, ultimately contributing to tumor growth control.

## Materials and Methods

### Bacterial Manipulation

Plasmid DNA was amplified in Escherichia coli (E. coli) DH5α for standard plasmids and in SW102 for Bacterial Artificial Chromosome (BAC) recombineering of GRAd genomes.

### Bacterial Transformation

Plasmids containing Fc single-chain (scFv-Fc, also referred to as SC) format, flanked by GRAd32 or GRAd25 E3 homology arms, were synthesized (Biofab). DH5α transformation was performed by heat shock with 100 ng plasmid DNA. After recovery in LB, transformed bacteria were plated on LB-Ampicillin agar and incubated overnight. A single colony was expanded in LB-Ampicillin, and plasmid DNA was extracted using the QIAGEN Plasmid Maxi Kit.

### Recombineering of GRAd BACs

Homologous recombination was performed in E. coli SW102 carrying BAC-GRAd with chloramphenicol resistance. Recombineering was induced by heat shock at 42°C to activate λ-red recombination. The first step involved positive selection via electroporation of an AmpR/SacB selection cassette into the target region, followed by plating on LB-Chloramphenicol-Ampicillin. In the second step, counter-selection was applied using LB-Chloramphenicol-Sucrose to replace the cassette with the desired modification.

### Cell Culture

The following cancer cell lines - ovarian (SKOV-3), breast (MCF-7), lung (NCI-H727, NCI-H1975, NCI-H1299, A549), melanoma (A375m), glioblastoma (U87MG) - and normal cell lines (MRC5, HUVEC) were obtained from ATCC. Human Amniotic Mesenchymal Stromal Cells were a kind gift from Dr. Ornella Parolini (Università Cattolica del Sacro Cuore, Rome). Glioblastoma patient-derived cell lines (GCh6 and GCh14) were established as previously described^40^. Cells were cultured in DMEM, RPMI, or CHANG Medium supplemented with 10% FBS, 2 mM L-glutamine, 1% Penicillin-Streptomycin, and, where required, heparin (0.1 mg/ml) and ECGS (0.03 mg/ml). Cells were maintained at 37°C in 5% CO₂, passaged at 90% confluence using TrypLE Select (Gibco), and subcultured at ratios of 1:2 to 1:10, depending on cell type. All cell lines were tested routinely for mycoplasma contamination through polymerase chain reaction (PCR).

### Virus Rescue, Amplification, and Purification

Linearized viral DNA (PmeI digestion) was transfected into A549 cells using Lipofectamine 2000. Transfected cells were cultured until cytopathic effect (CPE) appeared (7-10 days post-transfection). Virus-containing lysates (P0) were harvested by freeze-thaw cycles, clarified by centrifugation, and used for subsequent amplification. Further amplifications were carried out by infecting A549 cells at a controlled multiplicity of infection (MOI) of 100, up to passages P2/P3. Once an adequate viral yield was obtained, the GRAd lysates were purified using the Vivapure® AdenoPack 20 RT kit (Sartorius). This system enables column chromatography purification of up to 1.0 × 10¹² group C adenoviral particles from a 20 mL culture. We efficiently used this method to purify both GRAdC and GRAdB viruses.

### MTS Assay

Cell viability was determined using the CellTiter 96® AQueous Non-Radioactive Cell Proliferation Assay (Promega), which measures the bioreduction of the MTS tetrazolium compound to formazan by metabolically active cells. Cells were seeded in 96-well plates pre-coated with ECM matrix (Applied Cell Extracellular Matrix, abm) at optimized densities for each cell line to ensure growth for up to 8 days. Cells were infected with virus at multiplicities of infection (MOI) of 100. The assay was performed 3 and 8 days post-infection (dpi) by adding 20 μl of MTS reagent per well, incubating for 1 hour at 37 °C, and measuring absorbance at 490 nm using an EnSight plate reader. Results were expressed as a percentage of metabolic activity relative to uninfected controls (mock = 100%).

### Infectivity Assay

Viral infectivity was quantified by detecting hexon-positive cells through immunostaining. Cells were seeded in ECM-coated 96-well plates (2.1 × 10⁴ cells/well) and infected with serial MOIs (100, 10, 1) in triplicate. After 48 h, cells were fixed and permeabilized with cold methanol (−20 °C, 20 min). After blocking with 1% BSA in PBS, cells were incubated with a mouse anti-Hexon primary antibody (AbCAM BO25 AD 05, 1:400) followed by an AF647-conjugated goat anti-mouse secondary antibody (Invitrogen A21235, 1:500). Fluorescent signals were detected and quantified using a CTL ImmunoSpot reader, and the number of spots per well was used as a measure of viral replication efficiency.

### Productivity Assay

Viral productivity was determined by quantifying newly produced viral particles following infection at 100 MOI. Cells were seeded in 24-well ECM-coated plates at optimized densities for each cell line and infected with viruses at MOI = 100, in duplicate.

Cells were incubated at 37 °C, 5% CO₂, and harvested at 3 and 8 dpi. Cell lysates and supernatants were pooled, subjected to three freeze–thaw cycles, and clarified by centrifugation. Viral DNA was quantified by droplet digital PCR (ddPCR), and productivity was expressed as viral particles per cell (vp/cell) according to the formula:

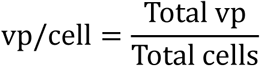

### GFP Expression Assay

To evaluate viral entry in cells non-permissive to viral replication, we used GRAd32 ΔE1 GFP, a replication-defective adenoviral vector expressing green fluorescent protein (GFP) under a constitutive promoter. MRC5 fibroblasts (non-permissive) and A549 cells (permissive control) were seeded in 6-well plates (1.0 × 10⁶ cells/well) and infected with the virus at MOIs of 100 and 1000. After 48 h, GFP expression was assessed by fluorescence microscopy, and representative images were acquired in both bright-field and GFP channels.

### GRAd-produced EV20 Antibody Expression Analysis

The expression of the EV20 scFv-Fc antibody was assessed in cells infected with GRAd32Fk25 EV20 and GRAd25 EV20 viruses. Human lung tumor cell lines (A549, NCI-H727, and NCI-H1975) and normal lung fibroblasts (MRC5) were seeded in 6-well plates at 1.0 × 10⁶ and 5.0 × 10⁵ cells/well, respectively, and infected with the indicated viruses at MOIs of 1, 10, 50, and 100. At 48 hours post-infection, supernatants were collected, clarified by centrifugation (4000 rpm), and analyzed by Western blotting analysis.

### GRAd-produced EV20 treatments

2.0 x 10^5^ A375m cells were seeded in 6-well plates in complete medium. After 24h, the cells were starved overnight in serum-free medium. The day after, the cells were washed with PBS and then pre-treated for 2h with adenoviral supernatants previously inactivated at 56°C for 30 min (1:20 diluted in serum-free medium), and with 10μg/ml of EV20 or heated EV20 as positive controls. After pre-treatments, cells were washed with PBS and stimulated for 10 minutes with 10 ng/ml of Human Neuregulin-1 (#5218, Cell Signaling) in serum-free medium. Cells were harvested and lysed in RIPA buffer (50mM Tris/HCl pH 7.6, 150mM NaCl, 1% NP-40, 0.5% Na-Deoxycholate, 0.1% SDS, 1mM EDTA pH 8) containing protease inhibitor cocktail (Sigma-Aldrich), phosphatase inhibitor cocktail (Roche, Thermo-Fisher), and Na_3_VO_4_ (Sigma-Aldrich). Lysates were clarified by centrifugation at 13000 rpm for 30 min at 4°C and subjected to Western blotting analysis.

### Western blotting analysis

For the analysis of GRAd-produced EV20 scFv-Fc antibody expression in cells infected with GRAd32Fk25 EV20 and GRAd25 EV20 viruses, 26 μL of each supernatant was mixed with NuPAGE™ LDS Sample Buffer (4×) and Sample Reducing Agent (10×) to a final volume of 40 μL, denatured at 99 °C for 10 min, and resolved on NuPAGE® 4–12% Bis-Tris gels. Proteins were transferred to nitrocellulose membranes, which were blocked for 1 h in 5% non-fat dry milk in PBS containing 0.1% Tween-20 (PBST). Membranes were then incubated for 1 h with an HRP-conjugated anti-human IgG secondary antibody (1:20000 dilution in PBST). After three washes, detection was performed using the Pierce™ ECL Western Blotting Substrate (Thermo Fisher Scientific), and chemiluminescent signals were acquired using a ChemiDoc Imaging System (Bio-Rad). To evaluate HER-3/Akt signaling cascade activation after GRAd-produced EV20 treatments, equal amounts of protein lysates (20 µg per sample), previously denatured at 95°C for 5 min, were subjected to SDS-PAGE electrophoresis and then electrotransferred to nitrocellulose membranes for Western blot analysis. Membranes were blocked for 1 h at RT with 5% non-fat dry milk in PBS with 0.1% Tween20. Membranes were then incubated overnight at 4°C with the following primary antibodies: anti Akt (#9272, Cell Signaling, Danvers, Massachusetts, USA), anti HER-3/ErbB3 (D22C5, Rabbit mAb, #12708s, Cell Signaling), anti Phospho-HER3/ErbB3 (Tyr1289, 21D3, Rabbit mAb, #4791s, Cell Signaling), anti Phospho-Akt (Ser473, D9E, Rabbit mAb, #4060, Cell Signaling), and anti-β-actin (mouse monoclonal; 1:40000, #A5441; Sigma-Aldrich). After three washes in PBS-0.1% Tween20, the membranes were hybridized with horseradish peroxidase (HRP)-conjugated secondary antibodies (rabbit or mouse; Biorad, CA, USA). Detection of signal bands was performed with Clarity Western ECL substrate (#1705061; Biorad). Images of membranes were acquired with a UvitecFire reader (Cambridge, UK) and analysed with Alliance Uvitec software (Cambridge, UK).

### Cytofluorimetric analysis

For the evaluation of HER-3 binding by GRAd-produced EV20 antibodies, 3.0 x 10^5^ A375m cells were collected and incubated for 30 min at 4°C with adenoviral supernatants inactivated at at 56°C for 30 min (1:2 diluted), and with 10 μg/mL of EV20 or heated EV20 (as positive control). Cells were washed with PBS and incubated for 30 min at 4°C in the dark with anti-human IgG AlexaFluor-488 conjugated secondary antibody (1:300, #A11013 Thermo-Fisher). For the evaluation of CD46 and CAR cell surface levels, 2-3 x 10^5^ cells (MRC5, HUVEC, A549, NCI-H1299, NCI-H1975, NCI-H727) were collected and incubated with anti-human CD46 (1:50; mouse monoclonal; #12239-MM05, Sino Biological) or with anti-human CAR (1:50; rabbit monoclonal; #10799-R271, Sino Biological) for 30 min at 4°C. Cells were then washed with PBS and incubated for 30 min at 4°C in the dark with anti-mouse or anti-rabbit AlexaFluor-488 conjugated secondary antibodies (1:300; #A11070, #A11017, Thermo-Fisher). Cells were analyzed using a FACS Canto II cytometer (BD Biosciences).

### Enzyme-linked immunosorbent assay (ELISA)

For the evaluation of binding affinity of GRAd-produced EV20 scFv-Fc antibodies, Maxisorp 96-well plates (Nunc, Thermo-Fisher) were coated with ECD (HER-3 C-terminal domain) (1µg/mL, 1:100 in PBS) overnight at 4°C. After blocking with 1% bovine serum albumin (BSA) in PBS for 1 h at RT, 100 µl of 1:2 or 1:10 diluted (in PBS) adenoviral supernatants (previously inactivated by heating at 56°C for 30 min) were added and incubated for 1 h at RT. For the detection, after three washes with PBS 0.05% Tween20, anti-human IgG-HRP (#A01070; Sigma-Aldrich) was added (1:5000) and incubated for 1 h at RT. After three washes, stabilized chromogen was added for at least 10 min in the dark before stopping the reaction with the addition of 1N H_2_SO_4_. The resulting color was finally read at 450nm with an ELISA plate reader (Tecan, Switzerland).

## Results

### Phylogenetic Classification of GRAd Viruses

To investigate the evolutionary relationships and classification of the gorilla-derived adenoviruses (GRAd), we performed phylogenetic analyses based on the hexon^41^, fiber, and polymerase genes. The goal was to determine the placement of GRAd viruses within established adenoviral species and assess their similarity to human adenoviruses (HAds) and simian adenoviruses (SAdVs).

The hexon-based phylogenetic tree (Figure 1A) shows that GRAd13, GRAd18, GRAd25 and GRAd54 (hereafter referred to as GRAdBs) clustered within group B adenoviruses, together with human adenovirus serotypes such as HAdV-35 and HAdB-11. In contrast, GRAd23 and GRAd32 (hereafter referred to as GRAdCs) grouped with group C adenoviruses, closely related to HAdV-5 and HAdV-6. Comparable results were observed with fiber-based (Figure 1B) and polymerase-based (Figure 1C) phylogenetic analyses.

**Figure 1.**
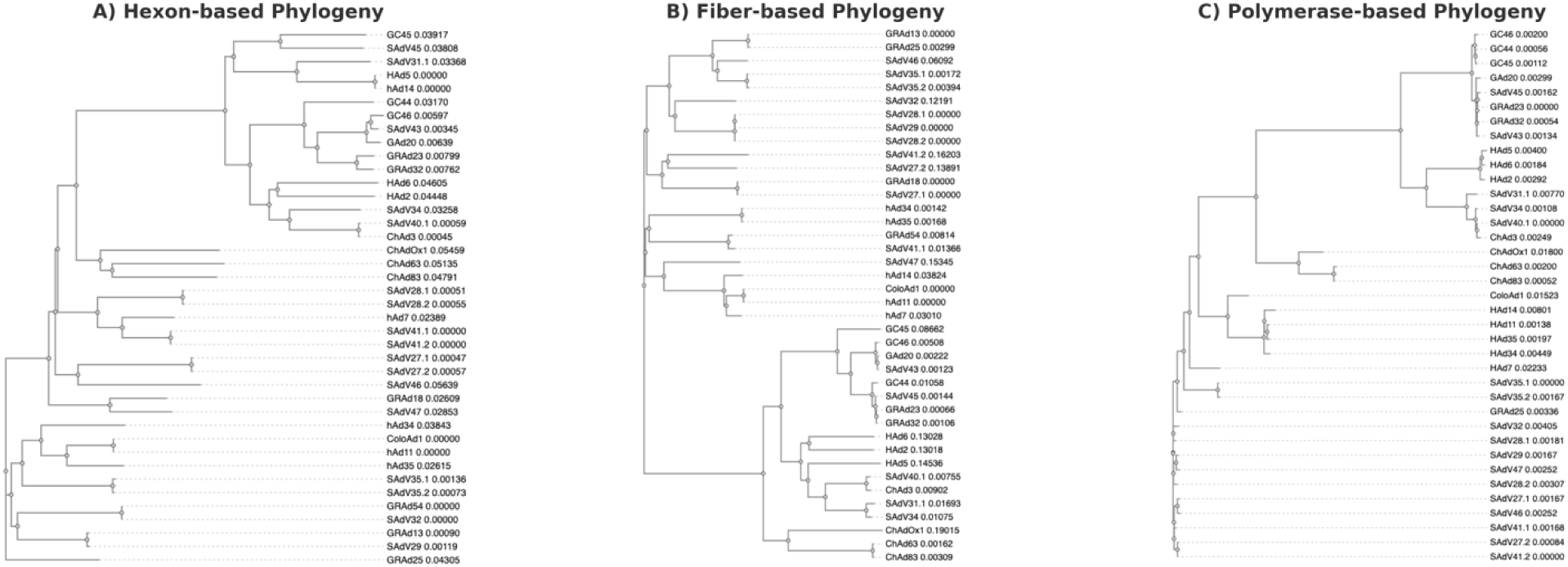
Phylogenetic classification of gorilla-derived adenoviruses from ReiThera. Phylogenetic trees were generated based on hexon (A), fiber (B), and polymerase (C) gene sequences to classify GRAd viruses within adenovirus species B and C. Multiple sequence alignments were performed using Clustal Omega, and maximum-likelihood phylogenies were inferred using MEGA11 with 1000 bootstrap replicates. GRAdB (13, 18, 25, 54) cluster within Group B adenoviruses, while GRAdC (23, 32) are closely related to Group C adenoviruses, confirming their evolutionary distinction. Bootstrap values (percentage of 1000 replicates) are indicated at branch points. Scale bars represent substitutions per site.

### Selective Replication and Cytotoxicity of GRAdCs

To evaluate the replication potential and oncolytic activity of GRAd vectors, we compared the ability of GRAdCs and GRAdBs to affect the metabolic activity of lung cancer and normal cells. HAd5 was included as a reference control. Cellular metabolic activity following infection was assessed by MTS assay (Figure 2A). GRAdCs selectively reduced the metabolic activity of the lung cancer cell lines A549 and NCI-H727, while exerting minimal to no effects on other lung cancer cells (NCI-H1975 and NCI-H1299) and normal cells (MRC5, HUVEC, and mesenchymal stromal cells). In contrast, GRAdBs and HAd5 displayed broader cytotoxic profiles, decreasing metabolic activity across both cancerous and non-cancerous cell lines.

**Figure 2.**
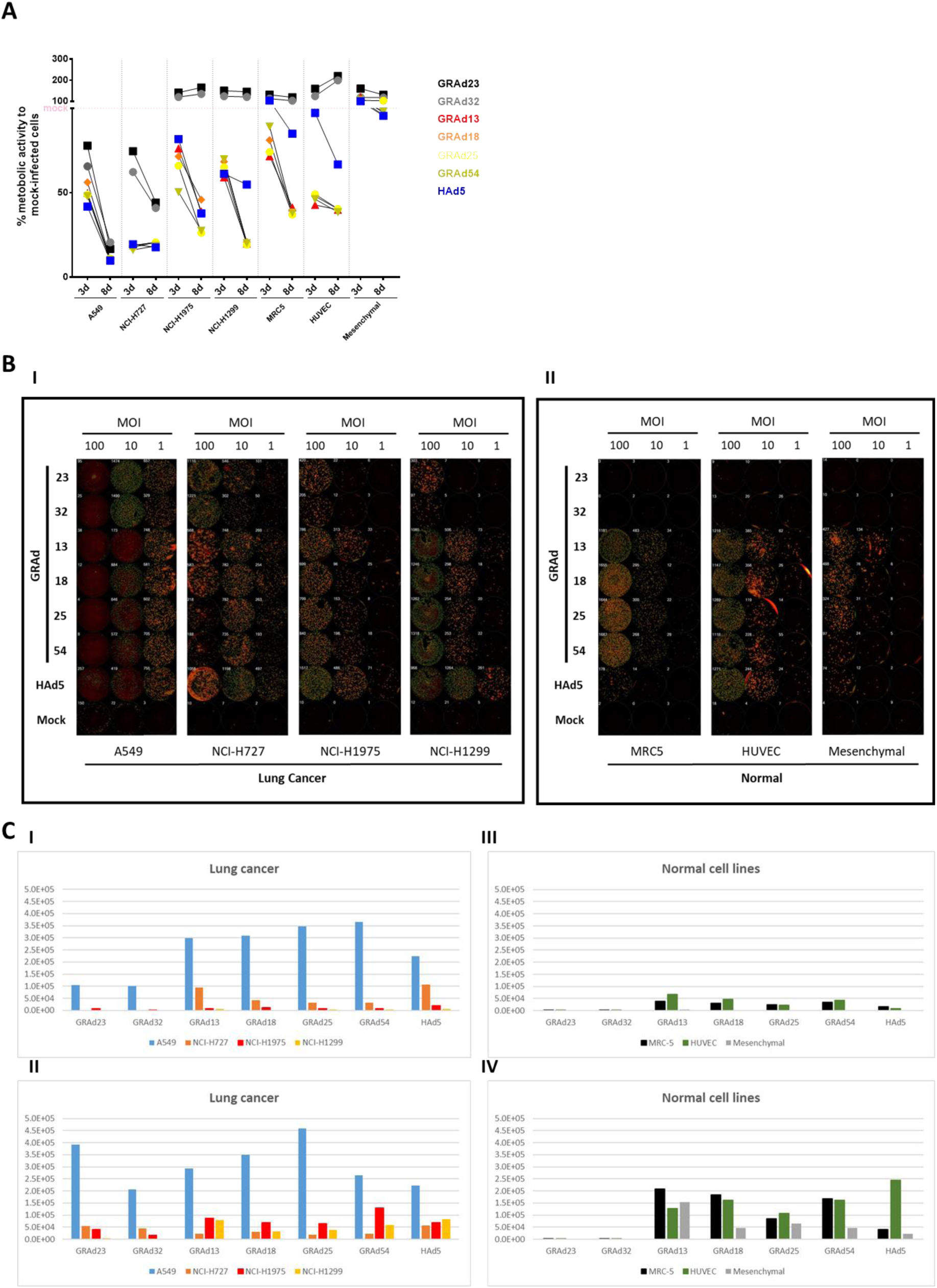
Comparative analysis of GRAdCs and GRAdBs replication, metabolic impact and productive infection in lung cancer and normal cells, in comparison to HAd5. (A) Metabolic activity of infected cells at 8 days post-infection measured via MTS assay. Lower metabolic activity indicates higher viral cytotoxicity. (B) Viral replication assessed through immunostaining for hexon protein at 48 hours post-infection. Higher numbers of hexon-positive spots indicate greater viral replication. (C) Viral genome production per infected cell at 3 days (I-III) and 8 days (II-IV) post-infection, measured via ddPCR, to assess productive infection.

Viral replication was next evaluated by quantifying hexon-positive cells via immunostaining at 48 hours post-infection (Figure 2B). GRAd23 and GRAd32 yielded high numbers of hexon-positive cells in A549 and NCI-H727, but replicated inefficiently in NCI-H1975 and NCI-H1299 (Figure 2Bi), as well as in normal cells (Figure 2Bii). Conversely, GRAdBs and HAd5 exhibited widespread replication in all tested cell lines, consistent with their broad replication profile (Figure 2B).

To further assess productive viral replication, nuclease-resistant viral genomes - reflecting encapsidated viral DNA - were quantified via ddPCR at 3 days (Figure 2C i, iii) and 8 days (Figure 2C ii, iv) post-infection. GRAdCs generated high viral genome yields in A549 and NCI-H727 cells, but remained restricted in other lung cancer cell lines and in normal cell lines. In contrast, GRAdBs and HAd5 exhibited robust genome amplification in all tested cell types, supporting their lack of selectivity. Together, these findings indicate that GRAdCs possess a more tumor-selective replication profile than GRAdBs and HAd5, efficiently targeting a subset of lung cancer cells while sparing normal cells.

### Modifying GRAd32 to improve tropism through chimeric Fiber engineering

GRAdCs showed strong selectivity for a subset of lung cancer cell lines but failed to replicate efficiently in other lung cancer models, limiting their therapeutic scope. We hypothesized that this restriction was driven, at least in part, by differences in receptor expression across cell types. While group C adenoviruses use the coxsackievirus and adenovirus receptor (CAR) for cell entry, group B adenoviruses typically use CD46, a receptor more uniformly expressed across many tumor types^42^. To demonstrate that GRAd25 utilizes CD46 as its primary entry receptor, A549 cells were infected with either GRAd32 or GRAd25 at a low MOI in the presence or absence of an anti-CD46 antibody. The cells were then monitored over three days for the development of cytopathic effect (CPE). While GRAd25 readily induced CPE in the absence of the anti-CD46 antibody, this effect was almost completely blocked when the virus was co-incubated with the antibody. Notably, at the specified time point and MOI, GRAd32 did not induce CPE either in the presence or absence of the antibody, consistent with the higher replication capacity of group B viruses compared to group C viruses in this cell line (Figure 3A). Given that HAd5-based oncolytic vectors have frequently been modified to broaden tropism by incorporating alternative fiber knob domains^43,44^, we sought to enhance GRAdCs infectivity by engineering a chimeric fiber containing the knob domain of a GRAdB.

**Figure 3.**
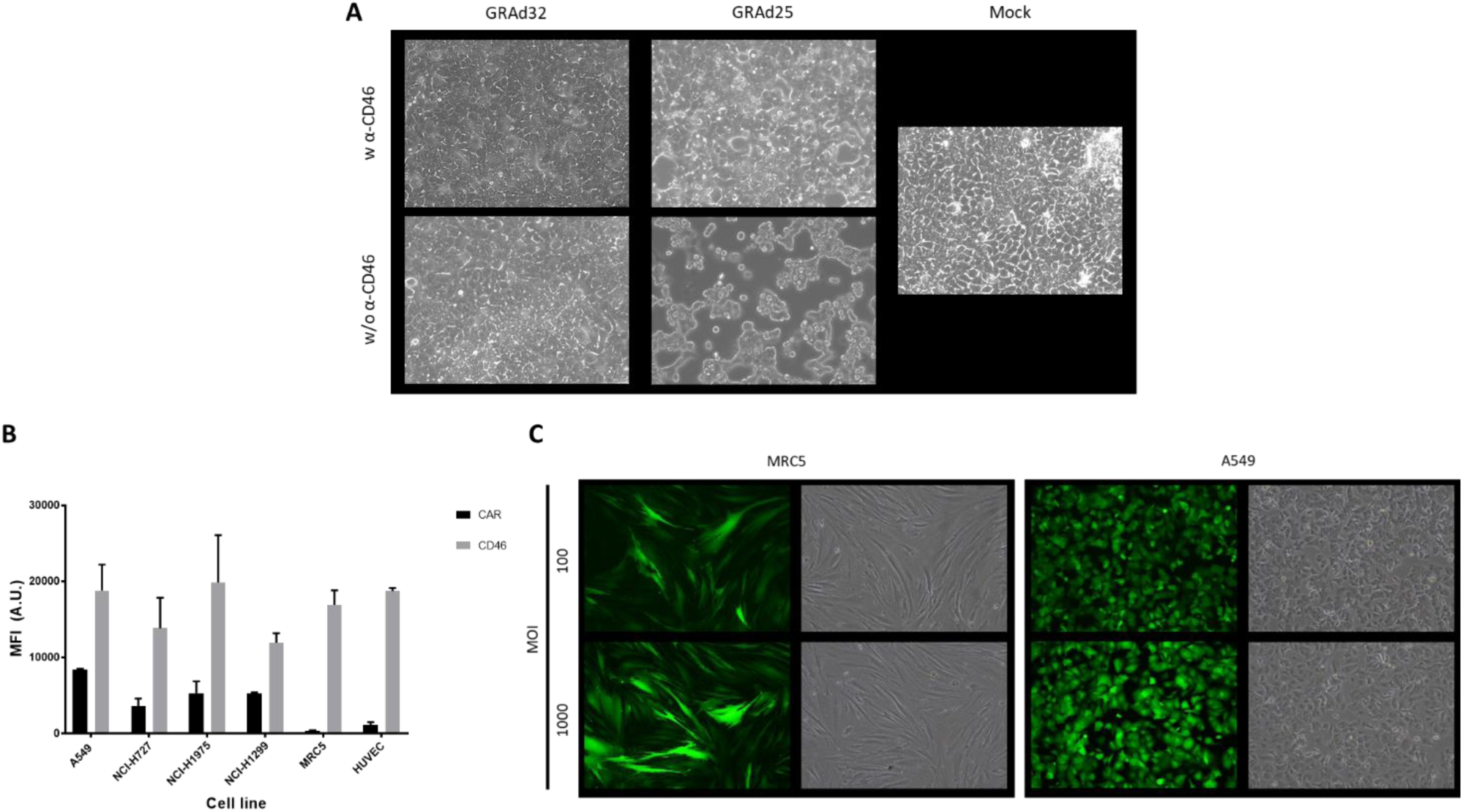
Side-by-side evaluation of receptor binding and cell entry mechanisms of GRAd32 and GRAd25. (A) A549 cells were infected with GRAd32 or GRAd25 at MOI 50 in the presence (w) or absence (w/o) of an anti-CD46 blocking antibody. Images were acquired 3 days post-infection. Competition with CD46 resulted in a delayed onset of cytopathic effect (CPE) exclusively in GRAd25-infected cells. (B) Flow cytometry analysis of surface CAR and CD46 expression levels in lung cancer (A549, NCI-H727, NCI-H1975, NCI-H1299) and normal (MRC5, HUVEC) cell lines. (C) Infection of MRC5 and A549 cells with a replication-defective GRAd32 GFP reporter virus. GFP expression was evaluated at 48 hours post-infection. The presence of GFP signal confirms that GRAd32 is capable of entering MRC5 cells despite its inability to replicate in this cell line.

To guide this strategy, we first evaluated CAR and CD46 expression in lung cancer cell lines (A549, NCI-H727, NCI-H1975, NCI-H1299) and normal cell lines (MRC5) by flow cytometry (Figure 3B). CAR expression was generally lower and more heterogeneous, whereas CD46 expression was more uniform across cells. Next, we tested whether GRAd32 could enter cells in which replication was not detected. To this end, A549 and MRC5 cells were infected with a replication-defective GRAd32 expressing GFP. At 48 hours post-infection, GFP expression was detected in both A549 and MRC5 cells, confirming that GRAd32 is capable of entering both cell types (Figure 3C), and suggesting that the replication restriction observed in MRC-5 likely reflects post-entry limitations. Based on these findings, we generated a chimeric GRAd32 in which the native fiber knob was replaced with that of GRAd25 while preserving the GRAd32 shaft and tail domains (Figure 4A). This construct, designated as GRAd32Fk25, was successfully rescued in A549 cells, and subsequently characterized in lung cancer and normal cell lines. To assess the impact of this modification on viral tropism, we measured metabolic activity by MTS assay at 8 days post-infection (Figure 4B). GRAd32Fk25 markedly reduced metabolic activity in tumor cells to levels comparable to GRAd25, while preserving the selectivity of GRAd32 by sparing normal cells.

**Figure 4.**
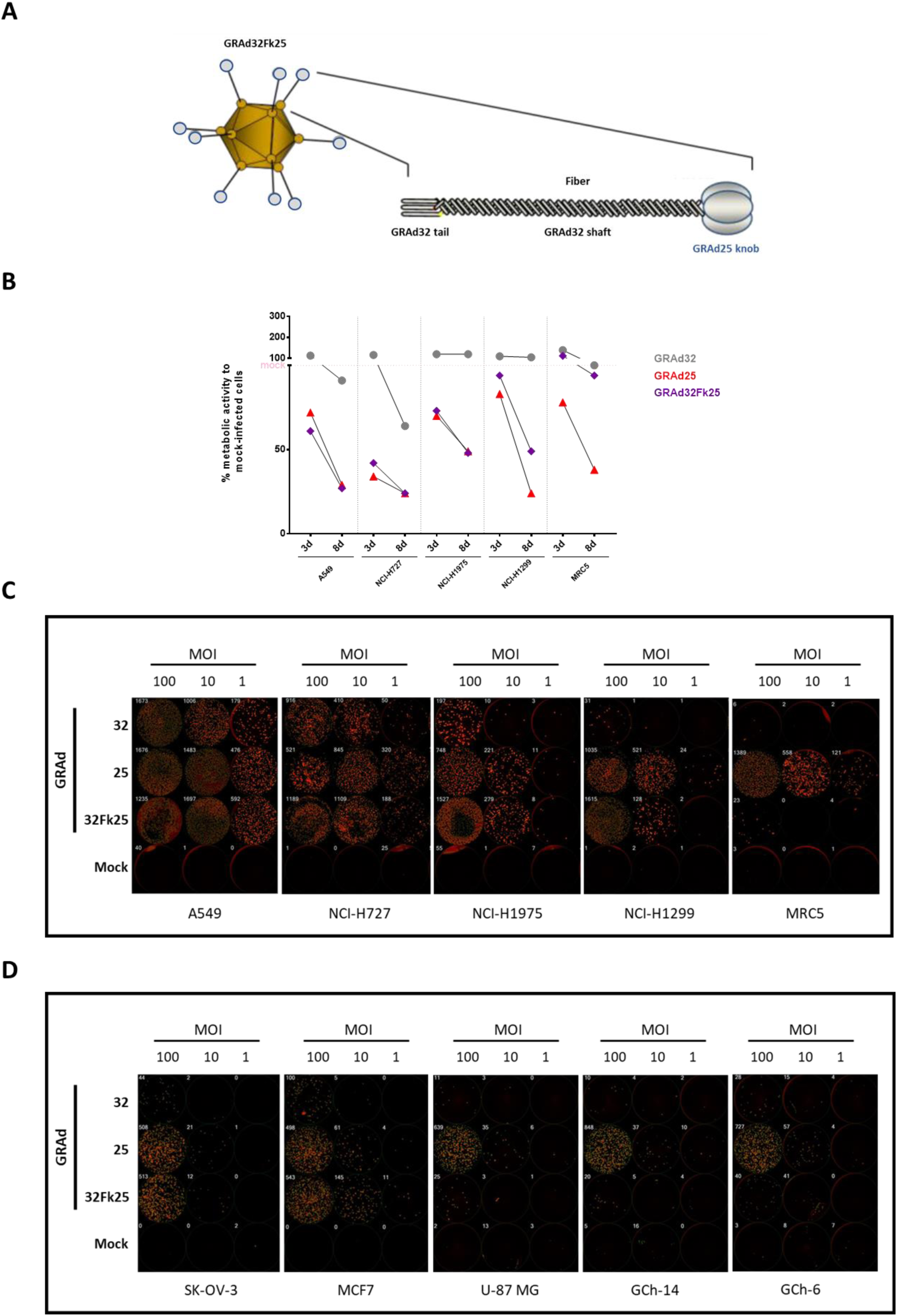
Engineering of GRAd32 with a chimeric Fiber to enhance tumor tropism. (A) Schematic representation of GRAd32Fk25. The chimeric virus retains the GRAd32 backbone, while its fiber displays a chimeric structure composed of the native tail and shaft domains of GRAd32 and the knob domain of GRAd25 (image adapted from ^55^). (B) Metabolic activity of lung cancer and normal cell lines infected with GRAd32, GRAd25, or the chimeric GRAd32Fk25 at MOI 100. The metabolic impact was measured at 3 and 8 days post-infection using the MTS assay. (C) Replication of GRAd32, GRAd25, and GRAd32Fk25 in lung cancer and normal cells at MOIs of 1, 10, and 100, assessed through hexon immunostaining at 48 hours post-infection. GRAd32Fk25 shows infection and replication efficiency similar to GRAd25 in tumor cells but remains restricted in normal cells. (D) Replication of replication-competent GRAd32, its retargeted derivative GRAd32Fk25, and GRAd25 was tested in ovarian (SK-OV-3), breast (MCF7), and glioblastoma cell lines (U-87 MG, GCh-14 and GCh-6). Red spots correspond to hexon-positive cells. The numbers displayed in each image indicate the total count of hexon-positive cells, serving as a quantitative proxy for viral replication efficiency.

To further evaluate infectivity and replication, cells were infected with GRAd32, GRAd25, or GRAd32Fk25 at MOIs of 1, 10, and 100, and viral replication was quantified by hexon immunostaining at 48 hours post-infection (Figure 4C). The number of hexon-positive cells was significantly increased in tumor cells infected with GRAd32Fk25 compared with parental GRAd32, consistent with enhanced infectivity. Importantly, while GRAd25 replicated robustly in both tumor and normal cells, GRAd32Fk25 retained a profile closer to GRAd32 in MRC5 normal cells, where it shows only limited replication.

Overall, these results show that replacement of the GRAd32 fiber knob with that of GRAd25 enhances viral entry and replication efficiency in tumor cells while maintaining a restricted replication profile in normal cells. This engineering strategy improves the oncolytic potential of GRAd32 without compromising its selectivity.

Next, we evaluated GRAd32Fk25 oncolytic potential across an additional panel of tumor cell lines, including breast (MCF-7), ovarian (SK-OV-3), and glioblastoma patient-derived cell lines (GCh6 and GCh14) (Figure 4D). In SK-OV-3 and MCF7 cells, GRAd32Fk25 achieved levels of replication comparable to GRAd25, substantially outperforming the parental GRAd32. On the other hand, in glioblastoma models, GRAd25 consistently produced numerous replication foci, whereas GRAd32 - either with its native fiber or with the GRAd25 fiber knob - produced none. Together, these data indicate that the retargeted fiber knob enables GRAd32Fk25 to acquire a GRAd25-like replication profile in cell lines in which GRAd32 replication is otherwise limited. Regarding glioblastoma, the lack of effect of GRAd32Fk25 may be explained by the fact that the underlying limitation lies in post-entry restrictions of group C GRAds, which are not overcome by the chimeric version of the virus.

### Oncolytic GRAd32Fk25 as a vector for Tumor-Site Antibody Therapy

We next engineered GRAd32Fk25 (and GRAd25 as a control) to produce a therapeutic monoclonal antibody within tumor cells. A single-chain Fc (scFv-Fc) version of the anti-HER-3 antibody EV20 (EV20 SC) was cloned into the E3 region of the viral genomes without an exogenous promoter (Figure 5A), and both viruses were rescued in A549 cells for further characterization.

**Figure 5.**
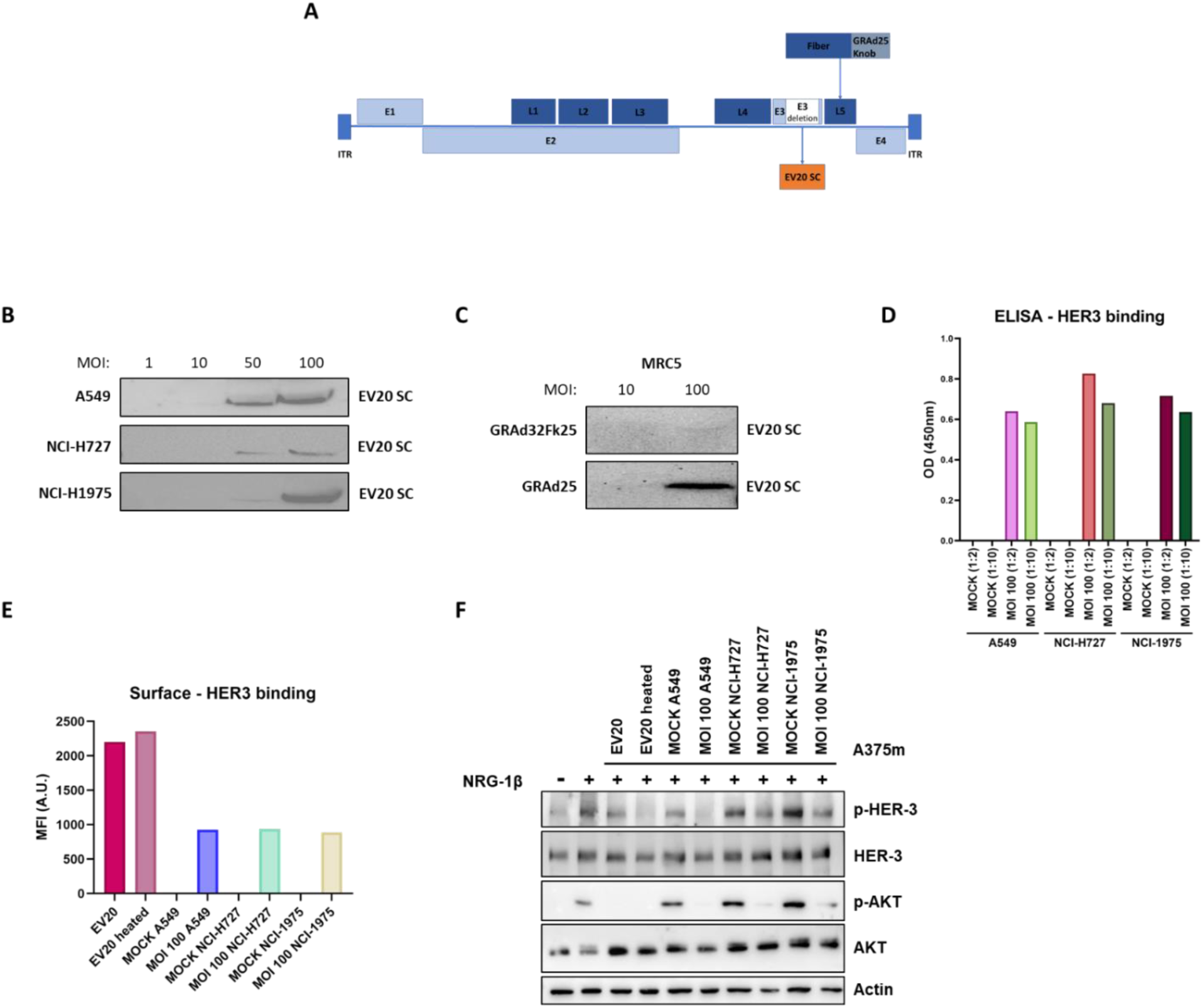
Expression and characterization of a therapeutic payload dependent on chimeric GRAd replication. (A) Schematic representation of the genetic structure of the GRAd32Fk25 EV20 SC vector. (B) A549, NCI-H727 or NCI-H1975 were infected with GRAd32Fk25 EV20 SC expressing a single-chain antibody targeting HER3, based on the EV20 sequence. Infection was performed at MOI 1-10-50-100, and supernatant-containing antibodies were separated on SDS-PAGE and blotted with an anti-human antibody. (C) MRC5 cells were infected either with GRAd32Fk25 expressing the EV20 SC, or with GRAd25 expressing the same EV20 SC at an MOI of 10 or 100. The supernatant-containing antibody were separated on SDS-PAGE and blotted with an anti-human antibody. (D) Affinity for HER3 (ECD) of supernatants containing GRAd-produced EV20 SC (1:2 or 1:10 diluted) derived from A549, NCI-H727, or NCI-H1975 infected with 100 MOI of GRAd32Fk25 EV20 SC or mock control. Affinity was evaluated through direct ELISA and absorbance values are expressed in OD (450nm) are reported. (E) Affinity for cell surface HER3 on A375m exposed to supernatants containing GRAd-produced EV20 SC derived from A549, NCI-H727, or NCI-H1975 infected with 100 MOI of GRAd32Fk25 EV20 SC or mock control. Affinity was evaluated by FACS and levels are expressed in arbitrary units (A.U.) of mean fluorescence intensity (M.F.I.). (F) Western blotting images for the evaluation of p-HER3, total HER3, p-Akt, total Akt, and actin (loading control) protein levels in A375m exposed to supernatants containing GRAd-produced EV20 SC (1:2 or 1:10 diluted) derived from A549, NCI-H727, or NCI-H1975 infected with 100 MOI of GRAd32Fk25 EV20 SC and treated or not with neuregulin-1β (NRG-1β). Equal amounts of protein samples were loaded per lane. All adenoviruses contained in the supernatants were heat-inactivated for 30 minutes at 56°C; EV20 is the positive control, EV20 heated is the positive control treated for 30 minutes at 56°C.

To confirm antibody production, lung cancer cells (A549, NCI-H727, and NCI-H1975) were infected with GRAd32Fk25 EV20 SC at MOIs of 1, 10, 50, and 100. At 48 hours post-infection, supernatants were collected and analyzed by Western blot using an anti-human IgG (Fc specific) antibody. EV20 SC was readily detected at higher MOIs in all tested tumor cell lines, indicating that the virus successfully expressed the therapeutic molecule in infected cancer cells (Figure 5B).

To assess the tumor selectivity of this approach, we compared the production of EV20 SC from GRAd32FkB25 with that from GRAd25 in normal MRC5 cells. As efficient antibody production depends on viral genome replication, the absence of significant expression in MRC5 cells would support replication-restricted expression. Western blot analysis showed strong antibody expression in MRC5 cells infected with GRAd25 EV20 SC, whereas only a faint band was detected for GRAd32Fk25 EV20 SC at MOI 100 (Figure 5C). This result is consistent with earlier observations that GRAd32Fk25 replicates only weakly in MRC5 cells, and further supports its selective profile.

Beyond confirming expression, we investigated whether the EV20 antibody produced by GRAd32Fk25 retained its functional capacity to inhibit HER3 receptor activation. *In vitro* binding assays confirmed that the antibody maintained its ability to bind HER-3 (Figure 5D). Flow cytometry analysis further demonstrated specific recognition of HER-3 on tumor cells (Figure 5E). Finally, we tested whether EV20 SC could inhibit PI3K/Akt signaling, a key downstream oncogenic pathway activated by HER-3. Western blot analysis of cancer cells treated with supernatant of lung cancer cells infected with GRAd32Fk25 EV20 SC revealed a clear reduction in phosphorylated Akt, indicating effective pathway inhibition (Figure 5F). Collectively, these data demonstrate that tumor cells infected with GRAd32Fk25 produce a functional EV20 SC antibody with comparable activity to purified full-length EV20, establishing GRAd32Fk25 EV20 SC as a potential tumor specific therapeutic with dual activity.

## Discussion

Our study highlights the potential of gorilla-derived adenoviruses (GRAd) as an oncolytic platform able to overcome key limitations of human adenoviruses, including preexisting immunity and off-target effects. GRAdCs displayed a highly restricted profile, entering only a limited number of cell types and replicating in even fewer, while GRAdBs exhibited broader cellular entry and replicated efficiently across essentially all permissive cells (a distinction not captured in Bots and Hoeben’s analysis^21^, which was limited to tumor cell lines). To combine these complementary features, and following an approach widely used in the adenovirus oncolytic field^43,44^, we generated GRAd32Fk25 by replacing the GRAd32 fiber knob with that of GRAd25, thereby achieving broader entry without loss of tumor selectivity. Indeed, using a panel of lung cancer and MRC5 as control of non tumoral cell lines we clearly demonstrated that replacement of the GRAd32 fiber knob with that of GRAd25 enhances viral entry and replication efficiency in tumor cells while maintaining a restricted replication profile in normal cells.

Despite the significant increase in specificity obtained with the chimera, we observed that GRAd32Fk25 displays a minimum level of replication in MRC-5 cells, indicating that the lack of replication observed with parental GRAd32 in this normal cell type is not due to an absolute post-entry block, but is at least partially attributable to restricted viral entry. In this context, the minimum increase observed for GRAd32Fk25 replication compared to parental GRAd32 in MRC-5 cells is consistent with enhanced entry mediated by the GRAd25 fiber knob. Nevertheless, GRAd32Fk25 does not reach the level of replication achieved by GRAdB vectors, indicating that increased fiber-mediated entry alone is insufficient to fully recapitulate the biological behavior of GRAdBs in normal cells. This difference likely reflects additional post-entry advantages intrinsic to GRAdBs, as well as the contribution of non-fiber capsid determinants to viral uptake and replication. In expanding our panel beyond the initial set, we confirmed that GRAd32Fk25 replicates in other tumor cell lines, including breast and ovarian cancer cells, while the addition of the GRAd25 fiber conferred no benefit in glioblastoma cells. This outcome is not unexpected: oncolytic strategies targeting glioblastoma with Ad5 typically exploit an RGD insertion in the native fiber rather than a group B knob^45,46^. Interestingly, our data further indicate that group B adenoviruses replicate more efficiently than group C adenoviruses in glioblastoma cells through mechanisms that likely extend beyond fiber-mediated entry. Once selective replication has been achieved - either through E1-based modifications or other genetic alterations - it would be of interest to investigate whether incorporation of an RGD-like motif into the group B fiber knob could further enhance oncolytic efficacy.

Overall, the enhanced replication of GRAd32Fk25 over GRAd32 demonstrated to be markedly more pronounced in several tumors than in normal cells, resulting in a favorable tumor-to-normal replication ratio. This quantitative selectivity is reminiscent of the therapeutic index described for the oncolytic adenovirus Enadenotucirev (ColoAd1)^10^, which similarly does not exhibit strictly tumor-exclusive replication but rather a markedly enhanced replicative fitness in malignant cells compared with normal tissues, outperforming conventional Ad5-based vectors. Notably, Enadenotucirev and its derivatives have been developed with systemic administration in mind and have been safely delivered intravenously in multiple clinical studies (reviewed in here^47^). Achieving a comparable route of administration with Ad5-based oncolytic vectors has proven considerably more challenging due to neutralizing interactions in blood, tropism for liver and spleen, and clearance by macrophages, as reviewed in here^48^, but eventually, intravenous delivery of Ad5 has been made possible by modification in either the hexon hypervariable regions (to reduce recognition by preexisting neutralizing antibodies as well as sequestration from serum coagulation Factor X^49,50^), the fiber^51^, as well by physical shielding approaches such as polymer or peptide coating, exemplified by PeptiCRAd platforms^52^.

Beyond these consideration, and while formal biodistribution studies will be required, it is noteworthy that a gorilla-derived adenovirus, GC46, has been reported to display a natural lung tropism with minimal hepatic sequestration following systemic delivery, in stark contrast to Ad5^19^. GRAd32 shares a high degree of sequence homology with GC46 at both the hexon and fiber levels, supporting the hypothesis that its in vivo biodistribution may similarly differ from that of Ad5. If GRAd32 were to recapitulate the biodistribution profile observed for GC46, it could represent an oncolytic vector intrinsically suited for intravenous administration, potentially obviating the need for deliberate hexon and/or fiber engineering or physical shielding strategies to avoid liver sequestration. In addition, GRAd32 is characterized by a naturally low seroprevalence of neutralizing antibodies^53^ in contrast to the non-human primates-derived group C adenoviruses described by Bots et al.^54^, further supporting GRAd32 candidacy for systemic delivery.

With the aim to enhance its antitumor activity, we engineered the GRAd32Fk25 cancer-selective chimeric adenovirus capable of producing a functionally active anti–HER3 therapeutic antibody. By inserting the sequence encoding a single-chain Fc (scFv-Fc) version of EV20 antibody into the E3 region, we successfully restricted EV20 SC production to tumor cells. In our case, the choice of the E3 region as an insertion site was guided by the non-cytotoxic and secreted nature of the antibody, as this locus tolerates additional expression cassettes without impairing viral replication in vitro, and has been widely used for arming oncolytic adenoviruses with secreted immunomodulators^7^. The tumor-selective expression is consistent with the intrinsic replication specificity of GRAd32Fk25 and provides a mechanistic basis for the synergistic interaction between HER-3 targeting and oncolytic virotherapy. This strategy, indeed, enables GRAd-infected tumor cells to act as biofactories for *in situ* EV20 SC production. Our data demonstrated that EV20 SC produced by tumor cells effectively modulates downstream HER-3 signaling in a similar fashion of full length therapeutic EV20. It is important to note that GRAd-mediated oncolysis is known to promote immunogenic cell death, while EV20 is capable of stimulating Fc receptor–dependent immune cell activation. Together, these mechanisms may establish a synergistic interplay between HER-3ErbB signaling blockade, virus-induced immunogenic cell death, and immune effector recruitment, ultimately amplifying antitumor activity. Future studies are demanding to evaluate this synergistic therapeutic effect *in vivo* using xenograft models in immunodeficient mice when using the SC EV20 format^38^, or, alternatively, syngeneic tumor models employing species-adapted transgenes.

## Conclusion

In summary, this study identified gorilla-derived adenoviruses as a versatile and biologically distinct oncolytic platform, and highlights GRAd32 as a uniquely tumor-selective group C adenovirus. By combining the intrinsic selectivity of GRAd32 with fiber retargeting from a group B adenovirus, we generated GRAd32Fk25, a chimeric vector that broadens tumor cell entry while preserving a favorable tumor-to-normal replication ratio, reminiscent of that described for Enadenotucirev (ColoAd1), yet within a group C backbone that retains the biological potency of this species while potentially circumventing the high seroprevalence and unfavorable biodistribution typically associated with Ad5. Beyond its selective oncolytic activity, GRAd32Fk25 represents also an effective biofactory for localized delivery of therapeutic moieties, as demonstrated here with the anti-HER3 antibody EV20. GRAd32Fk25-EV20 SC appears a promising anticancer agent that combines oncolytic potency with high local antibody production at the tumor site, resulting in improved efficacy and activation of the immune system. Importantly, the low seroprevalence of GRAd32 and its close homology to gorilla-derived adenoviruses with non-hepatic biodistribution profiles suggest that this platform may be intrinsically well suited for systemic administration, potentially without the need for extensive capsid engineering or physical shielding strategies. Together, these features position GRAd-based vectors as promising candidates for next-generation oncolytic virotherapies, warranting further investigation in appropriate *in vivo* models to fully define their safety, biodistribution, and therapeutic potential.

## Availability of data and materials

All data generated or analyzed during this study are included in this published article.

## Author contributions

RS, EC, AR and GS designed the study, supervised the project and wrote the paper.

RS, IC, VP and AP performed the experiments.

SC and GS provided funding.

## Acknowledgements

We thank Morena Pinti and Asia Pece for EV20 characterization.

We thank Ludovica Rossi for GRAd32Fk25 characterization.

## Funding

GS is supported by Fondazione-AIRC [Investigator Grant 2021-id 25696]

## Conflict of interests

GS is shareholder of Mediapharma s.r.l. and the other authors declare that they have no conflict of interest in this study.

AR, RS and SC are designed as inventors on a patent application filed based on the results of this work.

